# Non-linear microglial, inflammatory and oligodendrocyte dynamics across stages of Alzheimer’s disease

**DOI:** 10.1101/2025.03.25.645240

**Authors:** Aurélien M. Badina, Kelly Ceyzériat, Quentin Amossé, Alexandre Tresh, Laurene Abjean, Léa Guénat, Emilie Vauthey, Stergios Tsartsalis, Philippe Millet, Benjamin B. Tournier

## Abstract

Alzheimer’s disease (AD) is characterized by cognitive decline and neuropathological hallmarks including Aβ plaques and Tau tangles. Emerging evidence indicates oligodendrocyte (OL) dysfunction and demyelination also contribute to disease progression. Here, we analyzed OL markers and inflammatory gene expression in human hippocampal samples at early and late AD stages. In early AD, we observed OL and myelinating pathways downregulation, alongside microglial and astrocytic activation, as well as upregulated chemokine CCL2 and peripheral immune infiltration markers. In late stages, expression of OL-related genes and myelination pathways increase, with a higher NG2/MBP ratio, coinciding with decreased microglial coverage and peripheral immune markers. These findings indicate that early neuroinflammation may impair OL function, while attenuated immune activity in late AD allows partial OL recovery. This study provides insights into stage-specific inflammatory and myelin-related changes in AD, supporting the relevance of understanding oligodendrocyte dynamics and potential regenerative responses for future therapeutic strategies.

**Highlights:**

- Early AD: heightened microglial activation and peripheral infiltration.
- Late AD: reduced microglial presence and oligodendrocyte partial recovery.
- Neuroinflammation shifts toward remyelination-supporting conditions.

**Graphical abstract:** 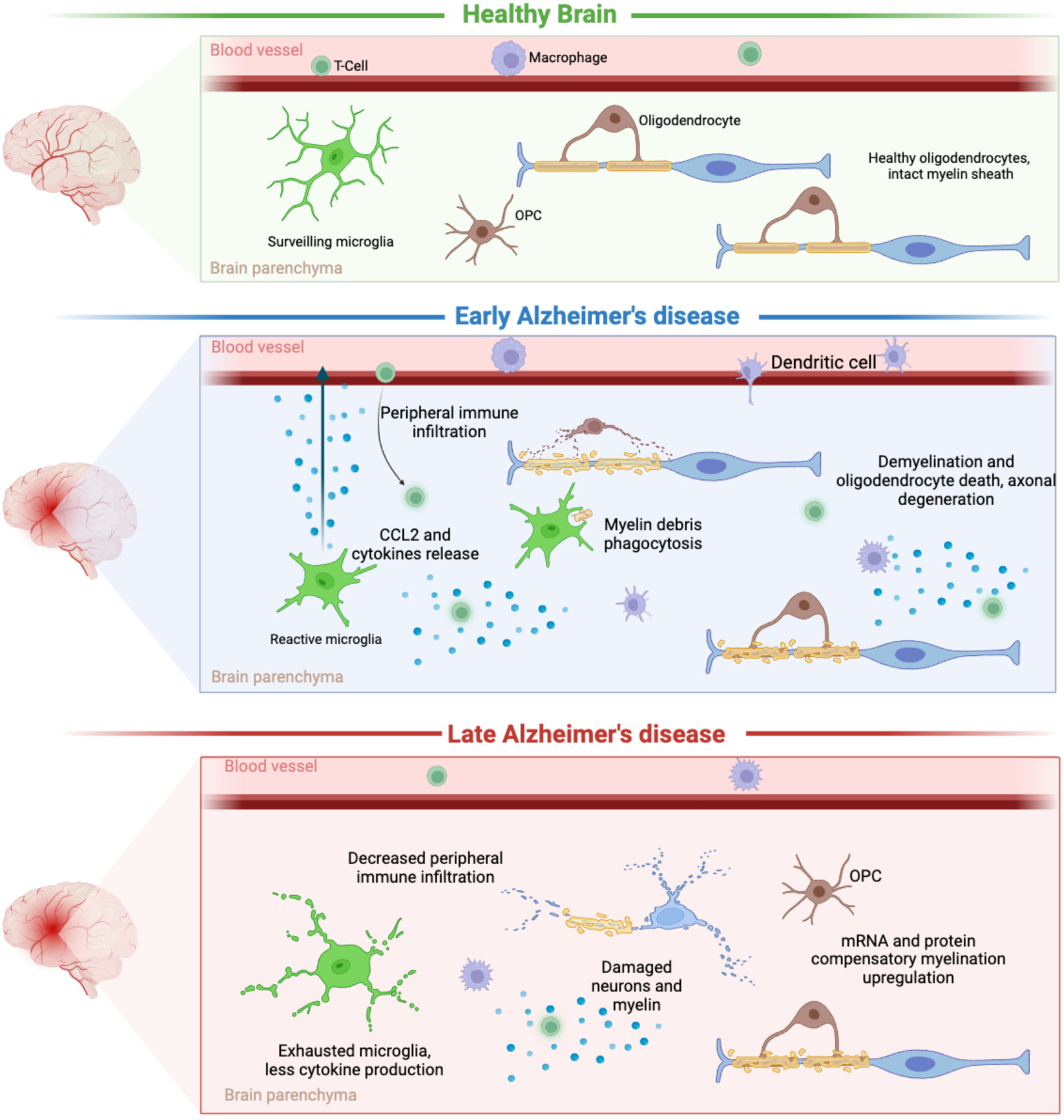

## Introduction

Alzheimer’s disease (AD) is the most common form of dementia and is linked to ageing, genetic and environmental factors and chronic neuroinflammation. Its sporadic form makes up to 95% of the cases and is linked to reduced amyloid β (Aβ) clearance in the central nervous system (CNS). The chronology of events leading to AD is debated but likely starts with an increased concentration of Aβ (Bloom, 2014). As Aβ clearance mechanisms are impaired, the peptides aggregate into plaques in the parenchyma. Aβ accumulation triggers glial reactivity, Tau tangle formation, inflammation, metabolic disruptions, and loss of neuronal support, all of which reinforce local toxicity and neuronal loss (Amro et al., 2021). The hippocampus is one of the first regions to be affected by neuronal loss, causing damages to memory formation mechanisms and spatial learning making it one of the key regions of interest (Rao et al., 2022). Traditionally, these pathologic mechanisms have been the primary focus of AD research. However, recent studies increasingly implicate disturbances in myelin integrity and oligodendrocyte (OL) function as critical contributors to disease progression (DeFlitch et al., 2022; Dong et al., 2018; Liu et al., 2025). Myelin, produced by OL, is essential for both electrical insulation of axons and for providing metabolic and trophic support to neurons. Disruptions in myelination not only compromise the rapid propagation of neuronal signals but also destabilize neural networks, potentially exacerbating memory deficits, learning challenges, mood disorders, and motor dysfunction (Boffa et al., 2022). Furthermore, oligodendrocyte precursor cells (OPCs) have the ability to proliferate and differentiate into mature myelinating OL after lesion or inflammation, making them an important factor in brain health (Ai et al., 2022; Franklin & Simons, 2022; Petiet et al., 2016). In AD, evidence suggests that OL dysfunction and demyelination occur early in the disease (Dong et al., 2018; Vanzulli et al., 2020), contributing to the progressive deterioration of neural connectivity. These suggest an important role for the myelinating unit in AD pathology and highlight the importance of understanding the mechanisms causing OL damage, and the potential for remyelination.

A key factor that may modulate OL health in AD is neuroinflammation. In the early stages of the disease, astrocytes and microglia become reactive in response to accumulating Aβ and Tau (Yu & Ye, 2015), releasing a host of pro-inflammatory cytokines and chemokines. Reactive glia produces pro-inflammatory cytokines, recruiting additional glial cells and facilitating the infiltration of peripheral immune cells, including macrophages, T-cells, monocytes, and Natural Killer (NK) cells to aid in pathogen clearance (Chen et al., 2023; Nevalainen et al., 2022; Rodríguez-Gómez et al., 2020). The infiltrated peripheral immune cells contribute to Aβ and Tau tangle clearance but also increase the risks of collateral toxicity and unintended damage. T-cells can engage with antigen-presenting cells, leading to demyelination. This effect is well documented in conditions such as multiple sclerosis, viral infections, and experimental autoimmune encephalomyelitis and is increasingly linked to AD (Bouhrara et al., 2018; Carmeli et al., 2013; Klotz et al., 2023). Notably, the monocyte chemoattractant protein 1 (also known as C-C motif chemokine ligand 2, or CCL2) is enough on its own to recruit peripheral immune cells and trigger demyelination, thereby contributing to these adverse effects (Errede et al., 2022; Kim & Perlman, 2005). This neuroinflammatory environment, while initially aimed at clearing pathological aggregates, impairs OL function and promote demyelination (Klotz et al., 2023; Larochelle et al., 2021).

Conversely, as AD progresses, microglial cells are documented to dystrophic phenotypes (Bisht et al., 2016; Frigerio et al., 2019; Hirbec et al., 2017; Keren-Shaul et al., 2017). These states involve morphological changes, such as process swelling, cytoplasmic deterioration, and cell fragmentation (Streit et al., 2009; Streit et al., 2020; Woollacott et al., 2020). They also include phenotypic shifts, with exhausted microglia displaying altered transcription factor production, reduced reactivity, and a decreased inflammatory phenotype compared to reactive microglia (Millet et al., 2024; Woollacott et al., 2020). A recent study suggests that interactions between microglia and Aβ or Tau may accelerate the transition from surveillant to senescent, potentially bypassing the reactive stage altogether (Shahidehpour et al., 2024).

In this study, we investigate the links between microglia, peripheral immune infiltration and OL functions in the human hippocampus. We analyze the dynamics of inflammation-related gene expression by quantification of their mRNA, and study how microglial activation and peripheral immune infiltration are linked to OL functions and pathways across AD stages. Microglial and macrophage presence is analyzed with IBA1 (coded by the *AIF1* gene), a marker that does not distinguish between resident microglia and differentiated infiltrated macrophages, marking a general macrophage presence fitting to test our hypothesis (Ito et al., 1998; Shahidehpour et al., 2025). Overall, we find a decreased IBA1 density in later stages, linked to a reduced immune infiltration and an upregulation of OL and myelination pathways. Through all this, we hypothesize that inflammatory conditions brought by reactive microglia and peripheral immune infiltration cause demyelination and OL death, which could be able to regenerate if those conditions subside.

## Material and methods

### Postmortem human samples

The neuroinflammation mRNA sequencing experiment was conducted on a cohort of fresh frozen samples originating from the Netherlands Brain Bank. The microglial staining experiment was conducted on a cohort of Formalin-Fixed Paraffin-Embedded (FFPE) samples originating from the Geneva Brain Bank. All experimental procedures were conducted in agreement with the Cantonal Commission for Research Ethics (CCER) of the Canton of Geneva.

### Fresh Frozen cohort for mRNA and protein analyses

The cohort was composed of 30 fresh frozen human hippocampal samples obtained from the Netherlands Brain Bank (Netherlands Institute for Neuroscience, Amsterdam, www.brainbank.nl). The subjects were distributed in three groups based on the Braak stages evaluated during autopsy, age- and sex-matched. The groups included 6 control subjects (CT, Braak 2-3), 12 early AD subjects (EAD, Braak 4) and 12 late AD (LAD, Braak 5-6). Detailed information are given in the supplemental data table 1. This cohort was used for mRNA and protein analyses.

### RNA extraction

Each fresh frozen sample was divided into two homogeneous halves. The first half was used for mRNA extraction by first being submerged in 400µl of TRIzol reagent (Thermo Fisher, 15596018). Samples were lysed by sonication and 100µl of chloroform per sample was added for phase separation. The samples were then centrifuged at 20’000g for 20 minutes at 4°C, and the supernatant was collected. mRNA was extracted using the RNeasy Mini Kit (Qiagen, 74104). Briefly, the samples were passed through filtrating columns to degrade proteins and other contaminants, washing away remaining material and keeping only genetic material following the company’s protocol. Deoxyribonuclease (DNase) was used during the extraction procedure to purify further the samples and keep only the RNA. The RNA was eluted in water and stored at −20°C until use. Quality control was done using electropherogram to ensure a sufficient mRNA quality.

### Protein extraction and quantification

The second half of the samples were used for protein extraction. They were suspended into a Triton X100 (Tx) lysis buffer concentrated at 50mM and pH 7.4, with 150mM NaCl and 1% Triton X100. Protease and phosphatase inhibitors were added to avoid protein degradation. The samples were sonicated, lysed and centrifuged at 20’000g for 20 minutes at 4°C. The supernatant containing the soluble/poorly aggregated proteins was extracted and kept at −80°C until use, and the pellet was resuspended in a Guanidine (Gua) lysis buffer containing 5M Guanidine in 50mM Tris HCl at pH 8, and protease and phosphatase inhibitors. The samples were then agitated on ice for 3 hours and centrifuged at 20’000g for 20 minutes at 4°C. The supernatant containing the highly aggregated proteins was collected and kept at −80°C until use. The following proteins were quantified by ELISA. On the triton extracted proteins (soluble, or poorly aggregated), we quantified Aβ40 and Aβ42 (Thermo Fisher, KHB3481 and KHB3544 respectively), pTau 231 (Thermo Fisher, KHB8051), CCL2 (Thermo Fisher, BMS281 Human MCP-1), IL21R (Life Technologies, EH265RB), NG2 (Thermo Fisher, EH337RB), MBP (LS Bio, LS-F4092). On the guanidine extracted proteins (insoluble, or highly aggregated forms), we quantified Aβ40, Aβ42 and pTau 231 using the same ELISA kits.

### RNA quantification

RNA quantification was performed based on the 770 mRNA probes present in the nanoString nCounter® Neuroinflammation Panel (XT-CSO-H-GLIAL-12). The procedure was done by the iGE3 Genomics Platform of the University of Geneva. The panel is composed of genes belonging to CNS cells such as astrocytes, microglia, oligodendrocytes, neurons as well as peripheral immune cells. The genes are involved in pathways such as cytokine signaling, innate and adaptative immune response and inflammatory signaling among others.

### Differential Gene Expression (DGE) analysis

Gene expression was normalized abased on the geometric mean of housekeeping genes and differential gene expression were calculated using nanoString’s nSolver (version 4.0.70). Volcano plots were produced based on the gene expression values with the EnhancedVolcano_1.24.0 package on R (version 4.4.2) (Blighe K, 2024).

### Cellular deconvolutions

Cellular deconvolutions were performed with the Expression Weighted Celltype Enrichment package (EWCE_1.14.0) (Skene & Grant, 2016) available on BioConductor (version 3.20).

### Pathway enrichment

Pathways enrichment was done with the clusterProfiler_4.14.6 package (Skene & Grant, 2016; Wu et al., 2021), and the gene symbols were converted to their Entrez Gene identifiers with the org.Hs.eg.db_3.20.0 package (M, 2025). Genes meeting the significance threshold were queried against GO databases (Molecular Function, Cellular Component, Biological Process, 2023 versions) to identify overrepresented pathways.

### Correlation analyses

The Pearson correlations were calculated with the stats_4.4.2 package (Wickham H, 2019) and plotted with the tidyverse_2.0.0 package.

### Gene set variation analysis (GSVA)

GSVA were done with the GSVA_2.0.5 package (Hänzelmann et al., 2013) and plotted with the tidyverse_2.0.0 package. Briefly, the method uses a Gaussian kernel to estimate the empirical cumulative distribution on a row (subject) basis. It computes a relative enrichment score, per subject, for each manually chosen gene set compared with all genes, then merges it with the original data to perform subsequent group statistics.

Formalin-Fixed Paraffin-Embedded (FFPE) hippocampus slices for IBA1 immunohistology The cohort was composed of 52 FFPE hippocampal slides obtained from the Geneva Brain Collection (Kovari et al., 2011). The subjects were distributed in groups based on their Braak stages evaluated during autopsy, age- and sex-matched. The groups were composed of 32 control (CT) subjects, 8 EAD subjects and 12 LAD subjects. This cohort was used for staining of microglial IBA1 marker. Detailed information are given in the supplemental data table 2.

### IBA1 immunohistochemistry and analysis

FFPE slides first underwent a deparaffinization battery of successive Xylene and degressive alcohol concentration baths and were submerged in 0.1% Glycine for 10 minutes at RT, then submerged in citric acid 0.01M at pH 6 for 20 minutes at 95°C, followed by successive baths of 0.1M PBS. Rabbit IBA1 primary antibody (Fujifilm, 019-19741) was diluted at 1:300 in 0.1M PBS, 0.3% Triton and 1% BSA and incubated overnight at 4°C under constant agitation. The slides were then rinsed in 0.1M PBS and incubated with secondary antibody HRP anti-rabbit (Dako, P044801-2) diluted at 1:100 in 0.1M PBS, 0.3% Triton and 1% BSA at RT for one hour. The slides were rinsed with 0.1M PBS and incubated with 3,3′-Diaminobenzidine (DAB) for 20 minutes at RT. After a last rinse with 0.1M PBS, the slides were then submerged in alcohol 60% and xylene and were then mounted with Entellan (Sigma-Aldrich, 1079600500) and imaged on a widefield scanner Zeiss Axioscan.Z1. IBA1 staining densities (covering percentages) were quantified using the QuPath software (version 0.4.3).

### Statistics

Protein levels across groups were analyzed with one-way ANOVA and Fisher’s LSD test. For the cellular deconvolutions, in these non-parametric tests, 10’000 random genes were generated to build a null distribution of enrichment scores for each cell type. To account for multiple testing across the 7 cell types, p-values were adjusted using the Benjamini–Hochberg false discovery rate (FDR) correction. Pathway enrichment was assessed using Fisher’s exact test, and p-values were adjusted for multiple comparisons using the Benjamini–Hochberg correction. The correlations between two markers were calculated with Pearson tests and linear regression line were fitted using the least-squares method. IBA1 densities across groups were calculated with Kruskal-Wallis and Dunn’s multiple comparison test. GSVA group statistics were calculated with a one-way ANOVA and post-hoc Tukey’s HSD test.

## Results

### I. Validation of AD groups composition and pathological markers

We validated the fresh frozen subject distribution in the control (CT), early Alzheimer’s disease (EAD), and late Alzheimer’s disease (LAD) groups by quantifying the poorly aggregated and highly aggregated forms of Aβ40, Aβ42 and pTau 231 (see Supplemental Fig 1). Consistent with literature, the pathological markers increased in our samples with disease progression. The fresh frozen and FFPE group distribution and composition are available in supplemental table 1 and 2 respectively.

### II. Inflammatory gene expression profiling and cell type enrichment

#### A. Differential gene expression across AD stages

We studied the inflammatory dynamics across the groups by quantifying the mRNA of 770 genes related to neuroinflammation and peripheral immune infiltration using nanoString’s nCounter neuroinflammation panel. Differential Gene Expression (DGE) visualized with volcano plots revealed different gene expression patterns between disease stages. Of the 770 possible genes, we observed 36 genes with dysregulated expression in EAD compared to CT, with 25 genes showing downregulated expression and 11 genes showing upregulated expression (Fig 1.A). This number increased to 278 genes showing dysregulated expression in LAD compared to CT, with 224 genes showing downregulated expression and 54 genes showing upregulated expression (Fig 1.B). LAD compared to EAD revealed 348 genes with dysregulated expression, with 261 genes showing downregulated expression and 87 genes showing upregulated expression (Fig 1.C). A Venn diagram visualization (Fig 1.D) revealed only three genes whose expression are significantly dysregulated throughout all the stages (see supplemental table 3), confirming the non-linear nature of inflammation-related dynamics in AD progression.

**Figure 1.**
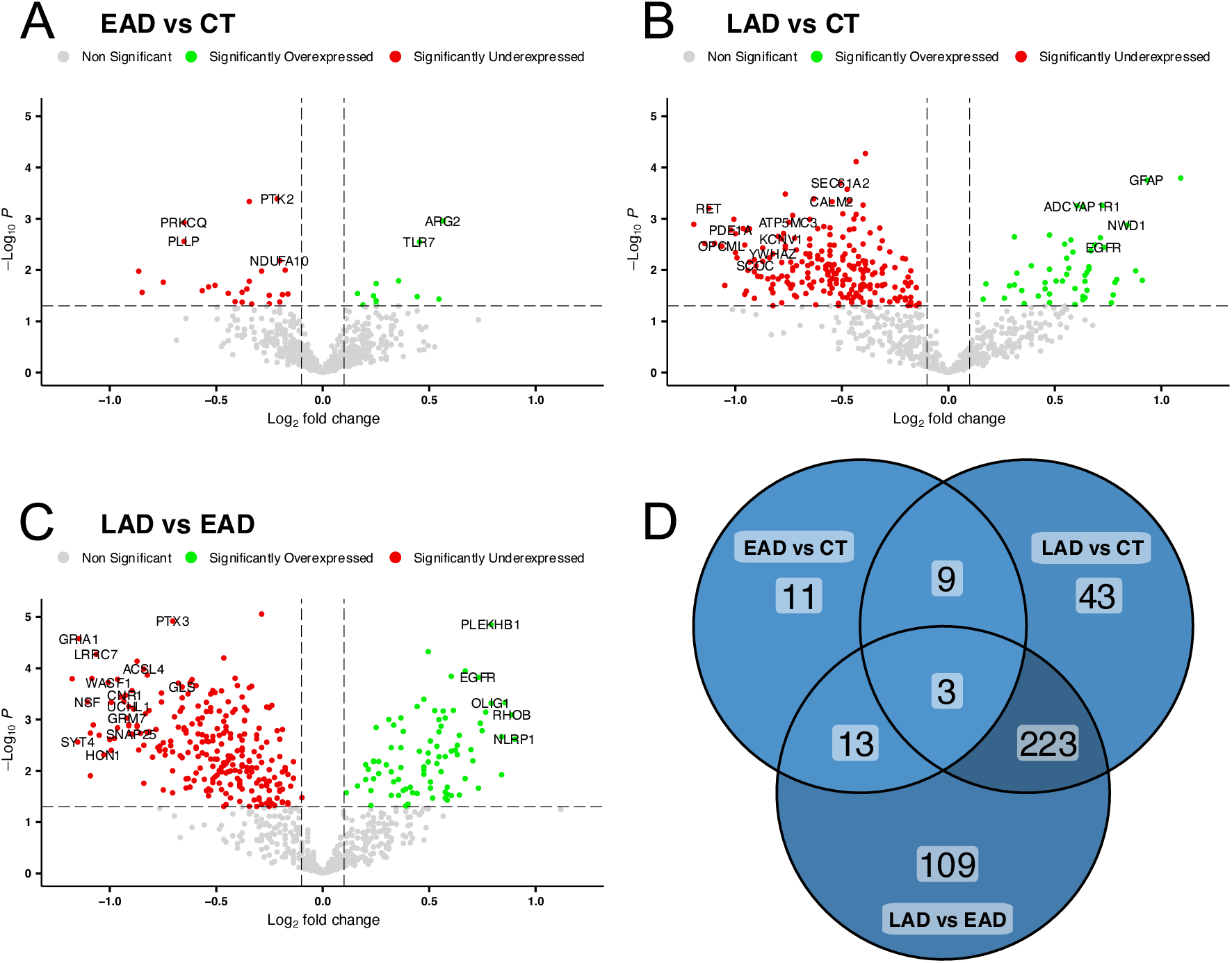
Volcano plots of Differential Gene Expression across stages. **A.** EAD versus CT: 36 genes showed dysregulated expression, with 25 downregulated and 11 upregulated. **B.** LAD versus CT: 278 genes showed dysregulated expression, with 224 downregulated and 54 upregulated. **C.** LAD versus EAD: 348 genes showed dysregulated expression, with 261 downregulated and 87 upregulated. **D.** Venn diagram of overlapping genes with significantly dysregulated expression. CT: Controls, EAD: Early AD, LAD: Late AD. Significance threshold is set at 0.05.

As we investigated differential gene expression and observed a non-linear inflammatory transcriptomic change with the disease’s progress, we then analyzed the pathways and functions mediated by the genes identified through those differentially expressed genes (DEGs).

#### B. Cell-type specific expression and pathway enrichment in early AD

Using cellular deconvolution and pathway enrichment (using the R packages EWCE and clusterProfiler, respectively), we identified cell types associated with the DEGs in EAD as compared to CT, and their related pathways with GO enrichment database.

In the early stages of AD, genes with downregulated expression are predominantly expressed by OLs (Fig 2.A, p-value<1e-4) and participate in pathways such as myelination and ensheathment of neurons (Fig 2.B). These genes include, among others, *MBP*, *MAG*, *GAL3ST1* and *PLP1*, all related to OLs. Conversely, genes with upregulated expression in EAD are expressed by astrocytes and microglial cells (Fig 2.C, p-values<0.01). They are associated with immune functions such as adaptive immune response, regulation of immune effector functions, chemokine production, and complement activation (Fig 2.D). These genes include *IL18*, *CFI*, *TLR7* and *C1S*.

**Figure 2.**
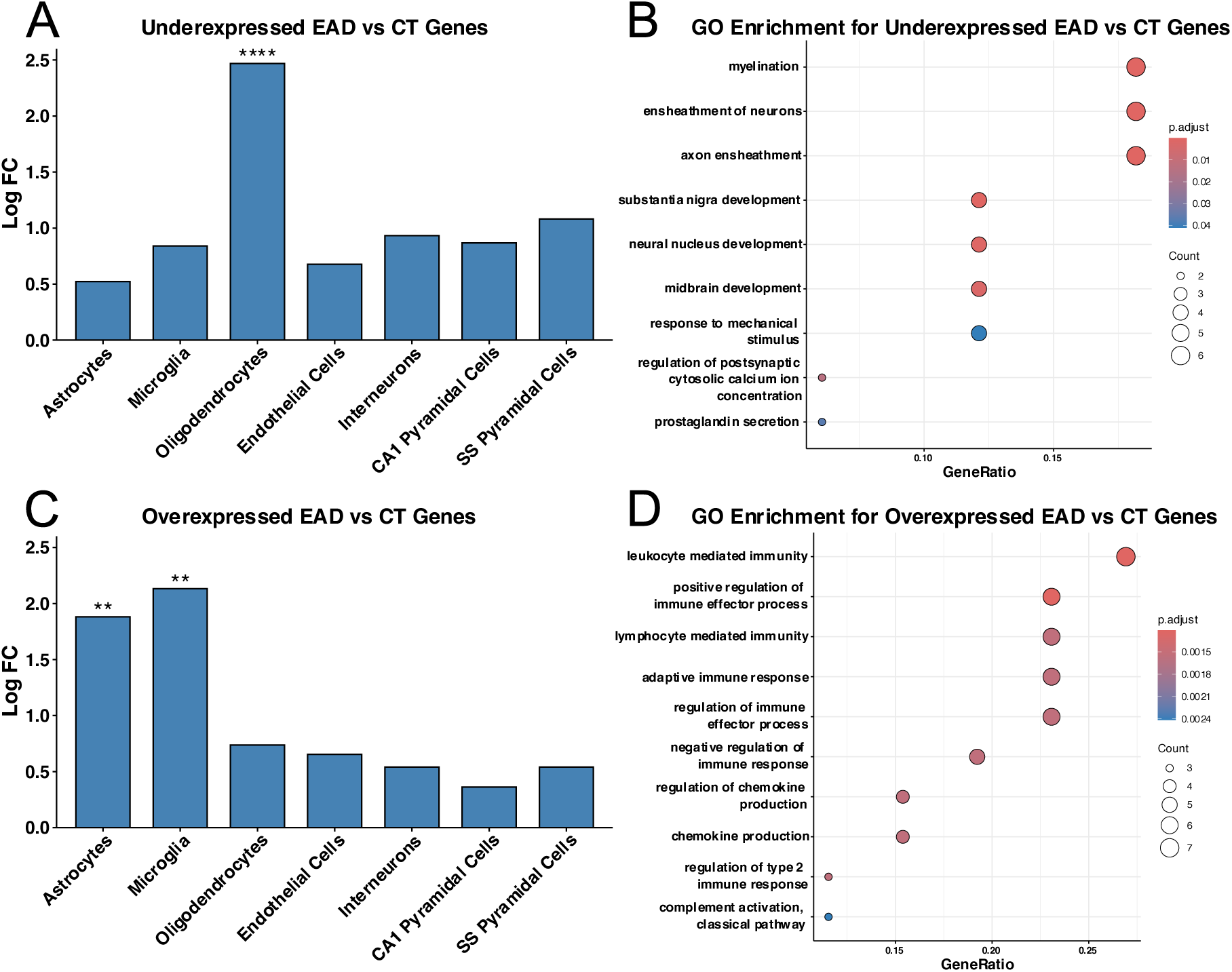
Cellular and pathway enrichment in gene changes in EAD versus CT. **A.** Cellular deconvolution shows that the genes with downregulated expression are expressed by OLs. **B.** Enrichment of genes identified in A reveals OL-related pathways such as myelination, ensheathment of neurons and axons. **C.** Cellular deconvolution shows that the genes with upregulated expression are expressed in astrocytes and microglia. **D.** Enrichment of genes identified in C reveals immune pathways such as regulation of chemokine production, immunity mediated by lymphocytes and leukocytes and complement activation. CT: Controls, EAD: Early AD. ** for p<0.01, **** for p<1e-4.

These results show that early AD is marked by increased astrocyte, microglial and inflammatory activity and occurs at the same time as decreased OLs function, myelination and neuron ensheathment.

#### C. Cell-type specific expression and pathway enrichment in late AD

Using the same cellular deconvolution and pathway enrichment methods, we then studied the gene showing dysregulated expression in LAD versus EAD to understand the dynamics in late stages of the disease. In LAD, genes with downregulated expression are predominantly neuronal (Fig 3.A, p-values<1e-4) and affect synaptic, axonogenesis and ion-transport pathways (Fig 3.B). They contain *SYN2*, *SNAP25*, *BDNF* and *GABRA3*. Genes with upregulated expression in LAD are expressed by astrocytes and OLs, but not microglia (Fig 3.C, p-values<1e-4). They are involved in pathways related to gliogenesis, glial cell differentiation, myelination and ensheathment of neurons (Fig 3.D). These genes include *OLIG1*, *OLIG2*, *SOX10*, *GAL3ST1* and *GFAP*.

**Figure 3.**
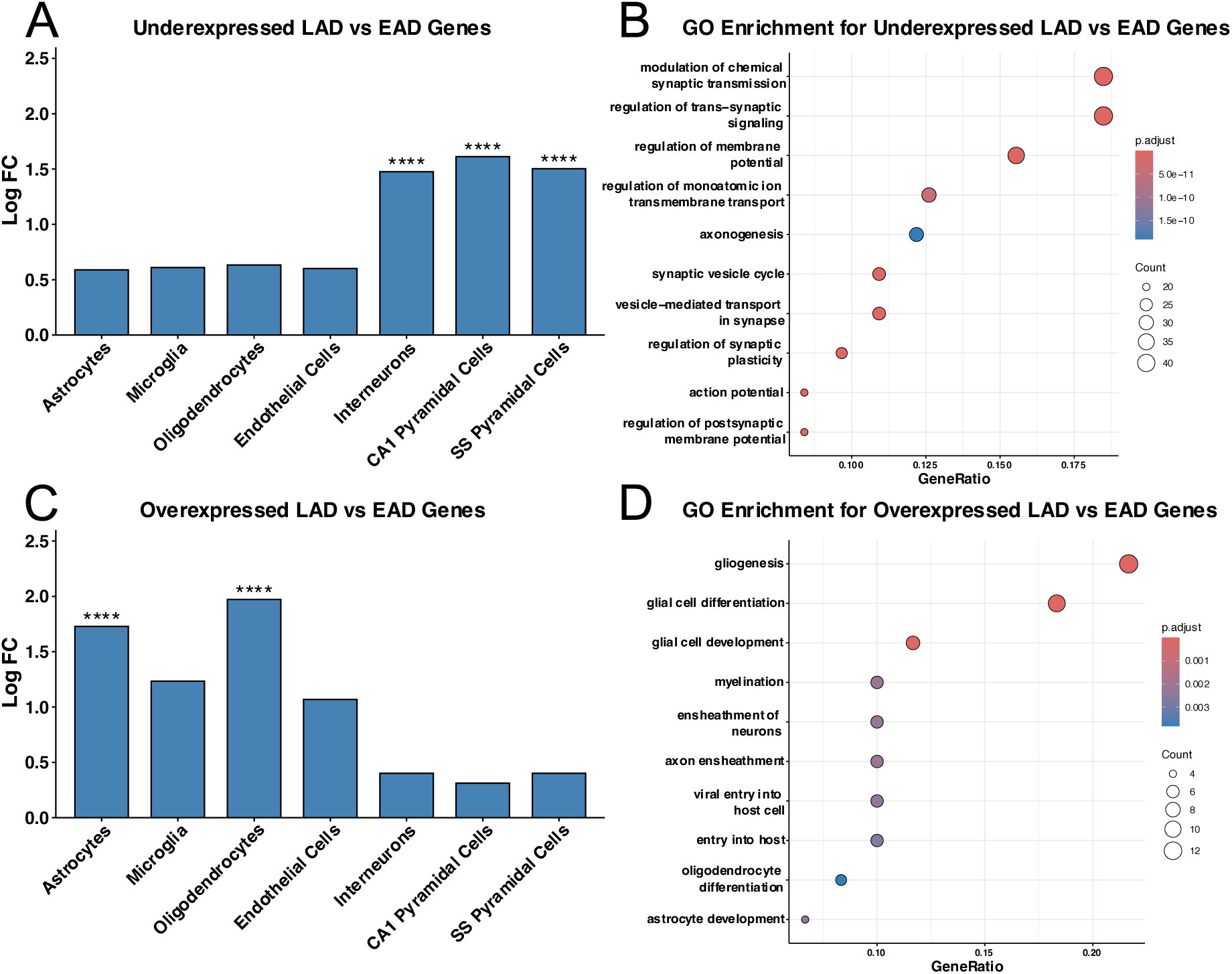
Cellular and pathway enrichment in gene changes in the progression from EAD to LAD. **A.** Cellular deconvolution shows that the genes with downregulated expression are expressed by different types of neurons. **B.** Enrichment of genes identified in A reveals neuronal pathways. **C.** Cellular deconvolution shows that the genes with upregulated expression are expressed in astrocytes and OLs but not microglia. **D.** Enrichment of genes identified in C reveals glial pathways such as OL development, gliogenesis, glial cell differentiation, myelination and axon ensheathment. EAD: Early AD, LAD: Late AD. **** for p<1e-4.

The shift from early to late AD is marked not only by further neuronal decline but also by a change in glial gene expression profile. Overall, we observed increased astrocyte activity through all stages, initially upregulated then constant microglial activity, as reflected by the impacted pathways, and initially downregulated then upregulated OL activity and myelin related genes and pathways. Moreover, the genes involved in OLs and myelin related pathways did not show differential expression between CT and LAD (see Supplemental Fig 2), indicating a transcriptomic return to control levels.

### III. Microglial activation and immune infiltration dynamics

#### A. Microglial presence by immunostaining

As microglial cells play a major role in inflammation and peripheral immune infiltration in the CNS, we then studied the non-linear microglial activity across the disease’s progression. To do so, we quantified the pan-microglia marker IBA1 (*AIF1*) on FFPE hippocampal slides of another cohort. Representative immunostaining scans of a CT subject (Fig 4.A), an EAD (Fig 4.B) and an LAD subject (Fig 4.C) are shown. The mean IBA1 density (percentage of covered area, Fig 4.D) over the scans increase from 4.21% (±2.9%) in the CT group to 10.92% (±5.5%) in the EAD group (p-value = 0.0015). IBA1 density then decreases back down to 5.27% (±4%) in the LAD group (p-value of 0.046). We observed no difference between the CT and the LAD group.

**Figure 4.**
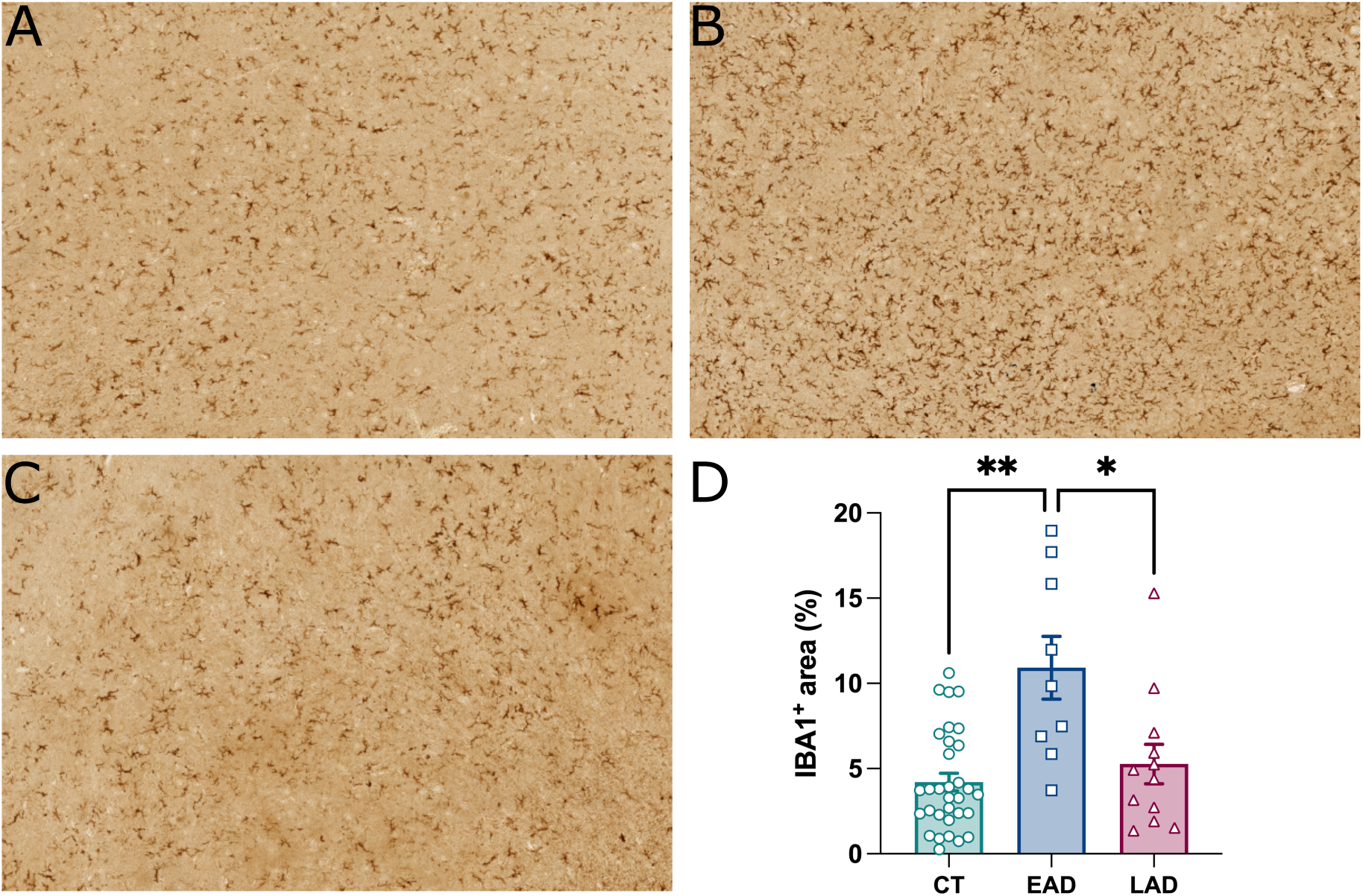
Microglial presence across stages of the disease. **A-C.** Representative DAB immunostaining of microglial marker IBA1 in a CT (**A**), EAD subject (**B**) and LAD subjects (**C**). **D.** Covering percentage (density) of IBA1 over hippocampal slides shows increased density from CT to EAD followed by a decrease in LAD with respective means of 4.21% (±2.92%), 10.92% (±5.51%) and 5.27% (±4%). Kruskal-Wallis and post-hoc Dunn’s multiple comparison test. CT *versus* EAD p-value=0.0015 and EAD *versus* LAD p-value=0.0462. CT: Controls, EAD: Early AD, LAD: Late AD.

These immunohistochemical analyses show that microglial presence, through the pan-macrophage marker IBA1, peaks in early AD and then decreases in later stages back to levels comparable to controls.

### B. mRNA correlations of inflammatory mediators with cellular markers

To further study the role of microglia on immune related functions, we analyzed the link between the expression of *AIF1* (IBA1’s coding gene) and infiltratory cytokine *CCL2*, showing a positive correlation of 0.65 (p-value = 0.0001, Fig 5.A). In turn, the expression of *CCL2* showed a positive correlation of 0.63 to Interleukin 21 Receptor (*IL21R,* p-value = 0.0002, Fig 5.B). *IL21R* is one of the specific markers expressed by peripheral immune cells and acts as an activation factor promoting the proliferation, differentiation, and maturation of T-cells, B-cells, and NK cells (Cagdas et al., 2021).

**Figure 5.**
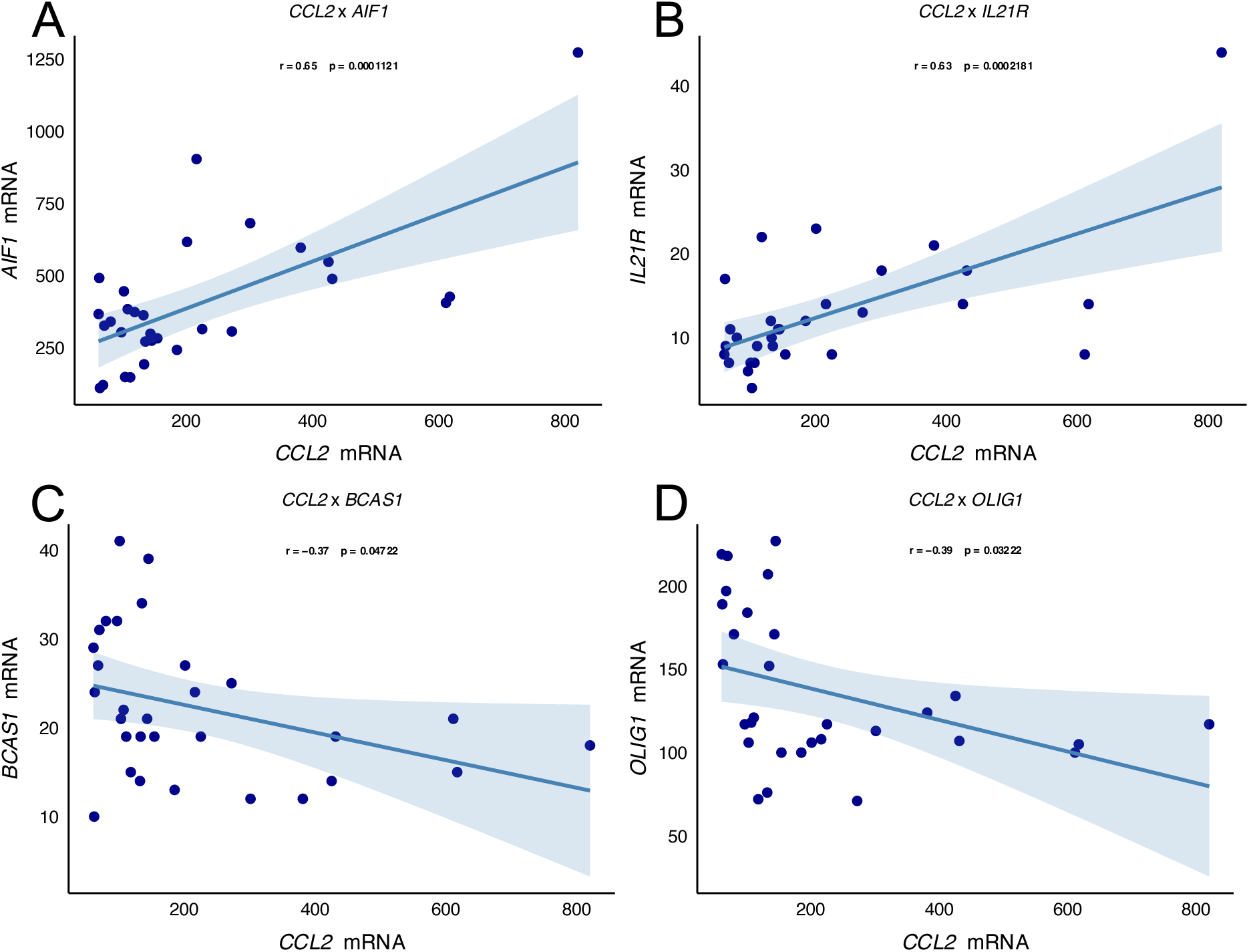
mRNA correlations between the expressions of *CCL2* and immune and OL genes. **A.** Positive correlation between infiltration-mediating CCL2 and microglial marker *AIF1*. Pearson correlation = 0.65 and p-value=0.0001. **B.** Positive correlation between infiltration-mediating *CCL2* and peripheral immune cells marker *IL21R*. Pearson correlation=0.63 and p-value=0.0002. **C.** Negative correlation between infiltration-mediating *CCL2* and regenerative OL marker *BCAS1*. Pearson correlation=-0.37 and p-value=0.047. **D.** Negative correlation between infiltration-mediating *CCL2* and OL transcription factor *OLIG1*. Pearson correlation=-0.39 and p-value=0.032.

Furthermore, the expression of *CCL2* was negatively correlated to Oligodendrocyte transcription factor 1 (*OLIG1)*, with a correlation of −0.39 (p-value = 0.032, Fig 5.C). *OLIG1* is a key player in OPC differentiation and the repair of demyelinated lesions (Zhao et al., 2019). It was involved in upregulated myelination pathways from EAD to LAD in our data (Fig 3.D). The expression of *CCL2* was also negatively correlated to another OL marker, Breast Carcinoma Amplified Sequence 1 (*BCAS1)* with a correlation of −0.37 (p-value = 0.047, Fig 5.D). *BCAS1* was recently studied for its importance in regeneration after inflammation and lesions in experimental cortical demyelination (Bergner et al., 2024).

These results show an inverse relationship between the expression of microglial inflammatory markers and those of OL regeneration and remyelination. To validate these dynamics and correlated expressions across stages observed on mRNA, in the next part, we quantified proteins involved in the same functions by ELISA.

### IV. Oligodendrocyte regeneration and remyelination responses

Protein quantification by ELISA confirmed a decrease quantity of infiltratory chemokine CCL2 between EAD and LAD, from 0.0727pg/µg (±0.057) to 0.03pg/µg (±0.016) (p-value = 0.026, Fig 6.A). The peripheral immune marker IL21R also showed a decrease between EAD and LAD, from 2.619pg/µg (±1.31) to 1.556pg/µg (±0.73) (p-value = 0.012, Fig 6.B). Conversely, the ratio of NG2 over MBP showed a strong increase from EAD to LAD with mean ratios of 10.32 (±3.62) to 91.8 (±93.6) (p-value = 0.009, Fig 6.C). NG2 (Neural/Glial Antigen 2) is a marker identifying OPCs and is rapidly upregulated following CNS injury (Polito & Reynolds, 2005). NG2^+^ cells are uniquely equipped to respond to CNS damage by differentiating into pre-myelinating OLs before maturing into myelinating OLs, a process critical for maintaining neural connectivity. On the other hand, Myelin Basic Protein (MBP) is a marker of mature myelin and one of the most abundant proteins in myelin sheaths (Dong et al., 2018; Vanzulli et al., 2020). Disruptions in MBP in AD are well known. These results indicate that NG2^+^ cells are more abundant in late stages than in early stages when compared to the overall myelin presence.

**Figure 6.**
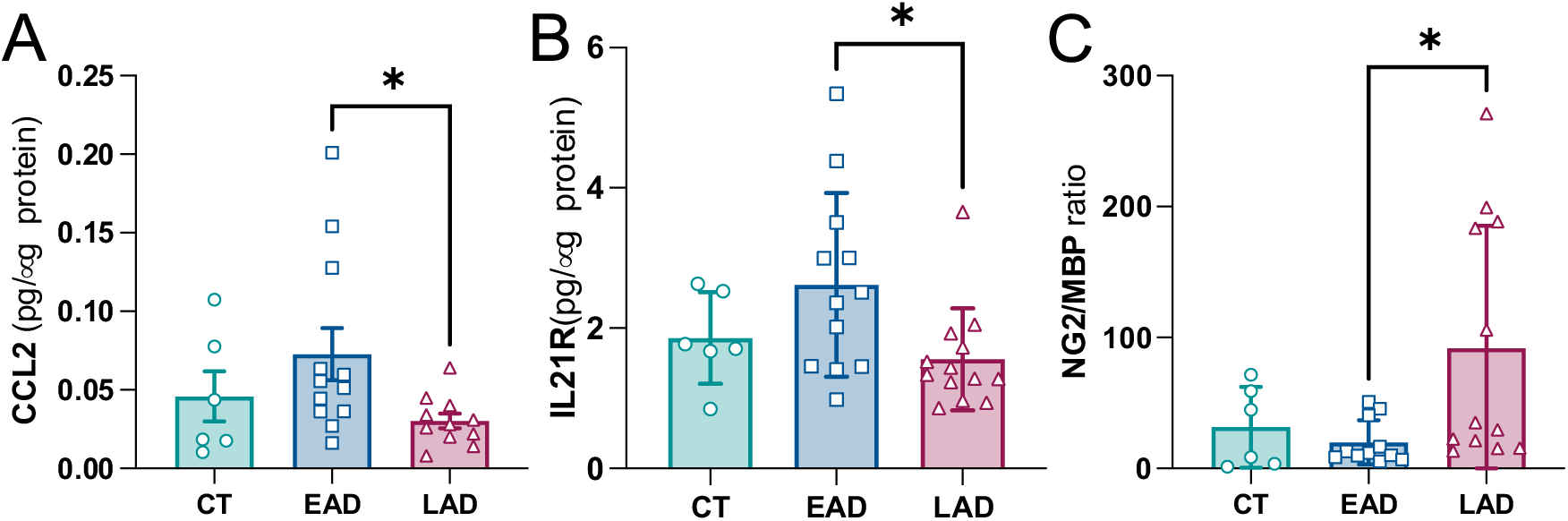
Protein dynamics of CCL2, IL21R and ratio of NG2/MBP across AD stages. **A.** Protein measurement of infiltration-mediating CCL2 across groups showing a decrease from EAD to LAD with mean concentrations of 0.0727pg/µg (±0.057) to 0.03pg/µg (±0.016). One-way ANOVA with post-hoc Fisher’s LSD test, p-value=0.026. **B.** Protein measurement of peripheral immune cells marker IL21R across groups showing a decrease from EAD to LAD with mean concentrations of 2.619pg/µg (±1.31) to 1.556pg/µg (±0.73). One-way ANOVA with post-hoc Fisher’s LSD test, p-value=0.0120. **C.** Ratio of OPC marker NG2 over mature myelin marker MBP across groups showing an increase from EAD to LAD with mean ratios of 10.32 (±3.62) to 91.8 (±93.6). One-way ANOVA with post-hoc Mann-Whitney test, p-value=0.0086. CT: Controls, EAD: Early AD, LAD: Late AD.

These protein results further confirm the mRNA expression patterns observed throughout the stages of the disease. As the disease progresses to late stages, we observed a decreased density of microglial marker IBA1, a decreased quantity of both infiltratory chemokine CCL2 and peripheral immune cells marker IL21R. Conversely, we observed an increased ratio of OPC over mature myelin from early AD to later stages, indicating a potential compensatory remyelination in the absence of microglial presence and peripheral immune infiltration.

Finally, to complement single-gene correlations, we performed gene set variation analysis (GSVA) to study the dynamics and correlations of functionally related gene sets (Fig 7.A) Using the gene data from the dysregulated pathways in the Fig. 2 and Fig. 3, we constructed a gene set involved in central and peripheral immunity and a gene set involved in OLs and myelination. To study further the hypothesis of changing microglial phenotypes, we also constructed a gene set involved in disease-associated microglia (DAM) (Silvin et al., 2022). Briefly, GSVA calculates a single enrichment score for the gene set in each sample compared to all other genes, reflecting relative up- or down-regulation of the gene set in the sample. Central and peripheral immune genes showed a strong increase in EAD maintained through LAD compared to CT (Fig 7.B). OLs and myelinating genes showed a decrease from CT to EAD, but no difference was observed between LAD and CT or EAD, representative of an intermediate stage (Fig 7.C). Genes associated with DAM showed an increase in LAD only (Fig 7.D). Central and peripheral immune gene enrichment scores negatively correlated with OLs and myelinating gene enrichment scores (Fig 7.E). Finally, the central and peripheral immune gene enrichment scores positively correlated with DAM gene enrichment scores (Fig 7.F).

**Figure 7.**
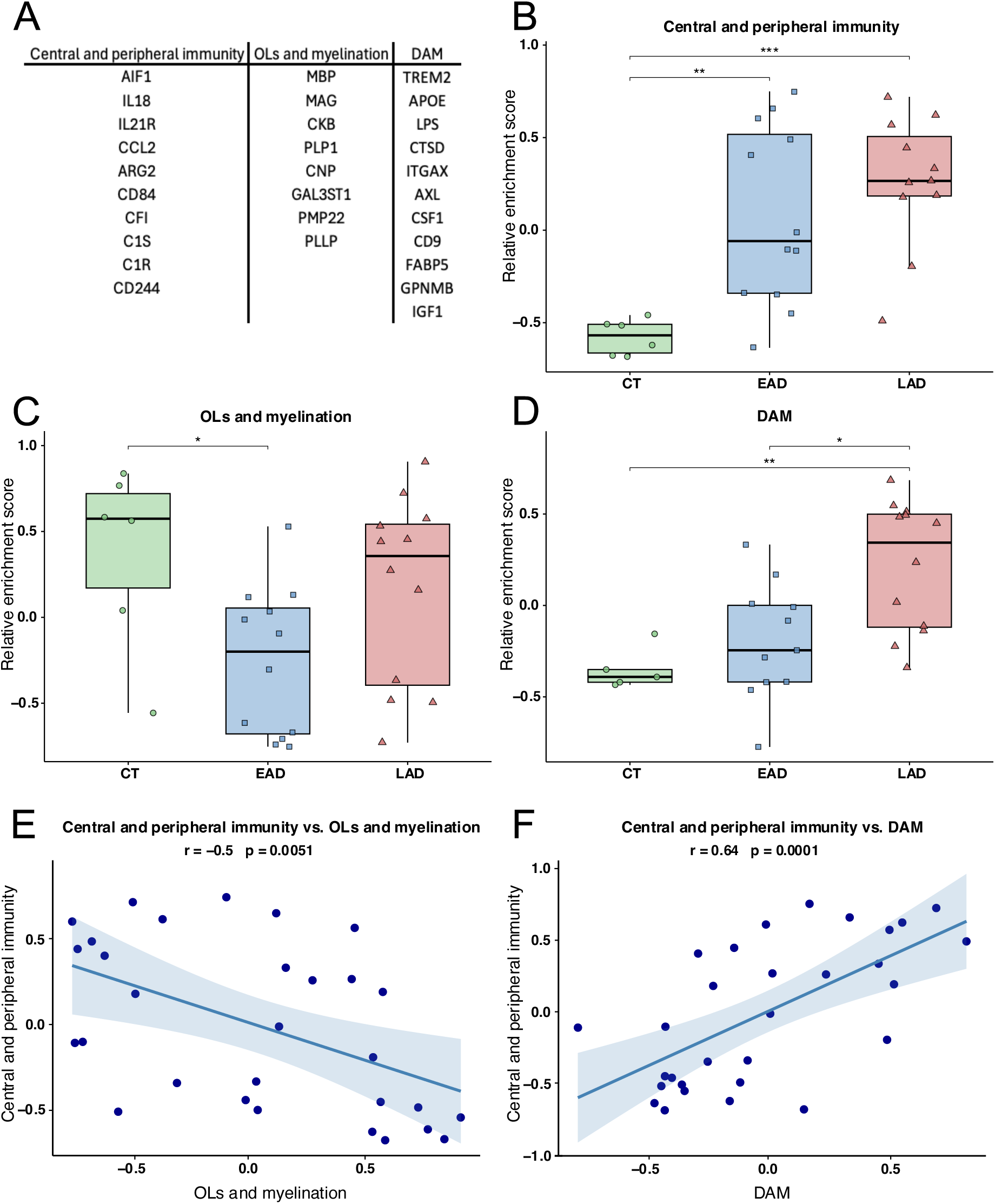
Gene sets variation analyses (GSVA) across stages and correlations of their relative enrichment scores. **A.** Gene names used in the tested sets. **B.** GSVA of genes involved in Central and peripheral immunity across stages showing an increase from CT to EAD (p-value=0.006) and LAD (p-value=0.0006). One-way ANOVA with post-hoc Tukey’s HSD test. **C.** GSVA of genes involved in OLs and myelination showing a decrease from CT to EAD (p-value=0.046) but no differences between LAD and CT or EAD. One-way ANOVA with post-hoc Tukey’s HSD test. **D.** GSVA of genes involved in disease-associated microglia (DAM) showing an increase in LAD compared to CT (p-value=0.006) and EAD (p-value=0.01). One-way ANOVA with post-hoc Tukey’s HSD test. **E.** Negative correlation between the expressions of the genes involved in central and peripheral immunity versus the expression of the genes involved in OLs and myelination. Pearson correlation=-0.5 and p-value=0.005. **F.** Positive correlation between the expressions of the genes involved in central and peripheral immunity versus the expression of the genes involved in DAM. Pearson correlation=0.64 and p-value=0.0001.

Through these results, we observed that the expression of immune genes is strongly increased in early stages of AD and are negatively correlated to physiological support functions mediated by OLs, while some infiltratory proteins such as CCL2 and IL21R fully recede as the pathology progresses. In late stages, the presence of IBA1 decreases compared to early stages, while the enrichment score for the DAM increases. This fully supports the idea of a changing microglial phenotype.

## Discussion

In this study, we examined how microglial activation, peripheral immune cell infiltration, and OL functions evolve over the course of AD. Our results showed a non-linear pattern of neuroinflammatory changes. Early stages were marked by heightened astrocytic and microglial reactivity, elevated cytokine production, and peripheral immune infiltration, whereas late stages displayed reduced microglial coverage and decreased levels of infiltratory chemokine (CCL2) and peripheral immune marker (IL21R). Importantly, we found a partial resurgence of OL and myelination-related gene expression in LAD, suggesting a potential remyelination attempt. These data underscore the complexity and stage-specific dynamics of inflammatory mechanisms in AD.

Early in the disease, our data revealed increased expression of genes linked to microglial and astrocyte reactivity, as well as upregulation of the peripheral immune cell infiltratory chemokine CCL2. This corresponds to a marked rise in IBA1 density in the hippocampus, indicating microglial activation (Ito et al., 1998). The elevated CCL2 and IL21R in EAD point to enhanced peripheral immune cell recruitment, consistent with the notion that microglia and astrocytes orchestrate the CNS immune environment. Indeed, CCL2, a critical chemokine for monocyte recruitment, is correlated with markers of both microglial and peripheral immune activity in our data. While CCL2 is known to induce neuroinflammation and demyelination in pathological contexts such as viral infections or encephalomyelitis (Dogan et al., 2008; Errede et al., 2022; Janssen et al., 2016; Kim & Perlman, 2005), it is less studied in AD. These findings align with prior literature indicating that microglia initially mount a pro-inflammatory response aimed at clearing Aβ and phosphorylated Tau (Castranio et al., 2022; Feng et al., 2020; Jäntti et al., 2022; Yu & Ye, 2015). However, in doing so, they produce cytokines and chemokines that recruit additional immune components, triggering collateral damage to surrounding OL and myelin structures. Prolonged inflammation does indeed lead to bystander damage to myelin and neurons (Hirbec et al., 2017; Park et al., 2021).

Based on these observations, we investigated whether heightened microglial activity correlates with changes in OL-related transcripts. In EAD, we found a downregulation of the expression of OL and myelin-related genes. Given the known relationship between chronic neuroinflammation and OL dysfunction or death, these results fit into a broader framework wherein inflammatory mediators and peripheral immune infiltration act as key stressors on the myelinating unit (Chatterjee et al., 2013; Klotz et al., 2023; Larochelle et al., 2021; Lee et al., 2015; Stohlman & Hinton, 2001). In accordance with literature, increased CCL2 levels correlated negatively with regenerative OL markers (*BCAS1*, *OLIG1*), confirming that microglial pro-inflammatory signals directly impede OL differentiation and myelin maintenance (Arfaei et al., 2024; Dogan et al., 2008; Errede et al., 2022). This effect is significant, as impaired myelination contributes to neural network instability and exacerbate cognitive deficits in AD, potentially accelerating disease’s progression and severity (Carmeli et al., 2013; Cheng et al., 2023). Taken together, our data strongly suggest that early-stage microglial activation, peripheral immune infiltration, and heightened cytokine production converge to promote myelin damage and reduce OL resilience.

Interestingly, as the disease progresses to late stages, microglial coverage, CCL2, and IL21R levels diminish, suggesting less peripheral immune infiltration. Although microglial cells remain present, our data propose a shift from a reactive phenotype to a dystrophic state. In accordance with literature, over time and as amyloid and Tau pathologies evolve, microglia may shift into a more dystrophic, exhausted or senescent phenotype as previously observed (Bisht et al., 2016; Keren-Shaul et al., 2017; Streit et al., 2020). This transition may reduce the intensity of pro-inflammatory and infiltratory signaling, thereby alleviating some of the pressure on OL lineage cells and partially permitting remyelination efforts. Such a change aligns with other reports that microglia lose their functional capacity to clear pathological proteins and dysfunctional over time (Keren-Shaul et al., 2017; Millet et al., 2024; Streit et al., 2020). Another explanation could simply be the death of microglial cells after chronic exposure to Aβ (Baik et al., 2016; Qiu et al., 2023). Both could concur, as our dataset shows the emergence of the DAM phenotype in late stages of the disease only. Coupled with the decrease of IBA1 density in late stages, microglia could both die or enter dysfunctional phenotypes due to chronic exposure to toxic elements and metabolic stress. Such dysfunctional states could prevent it from fulfilling some inflammatory roles, peripheral immune infiltration in our case. Interestingly, the central and peripheral immune gene set was highly correlated to the DAM gene set. This could suggest that while the peripheral immune infiltration decrease, local toxicity remains but shifts to being mediated by DAM and dysfunctional microglia. An inflammatory attenuation may create a window for remyelination, a concept previously proposed in Parkinson’s disease (Worlitzer et al., 2012). In parallel with these changes in microglial activation and reduced inflammatory signals, in our data, OL genes and myelination-related pathways partially re-emerged in LAD, accompanied by an increased ratio of the OPC marker NG2 over the mature myelin marker MBP. The analysis of the more complete gene set involving OLs and myelinating genes confirmed the decreased enrichment in early stages, and late stages present an intermediate state between controls and early stages of the disease. This is consistent with the fact that no healing has been reported in advanced forms of AD. However, it suggests an attempt by OPCs to differentiate into mature OLs and initiate remyelination, matching with studies in other neurodegenerative and demyelinating diseases, where prolonged inflammation eventually subsides or shifts phenotype, allowing a limited window for OL regeneration (Ai et al., 2022; Bergner et al., 2024; Petiet et al., 2016). Nonetheless, neuronal genes and pathways remained suppressed, showing that this remyelination attempt does not rescue neuronal markers expression.

As such, while these findings open perspectives for understanding dynamic inflammatory states and their consequences in AD, important questions remain, and several limitations need to be addressed. First, although we observed correlations among the expression of microglial markers, chemokines, and OL genes, causality cannot be fully established from these data alone. Future studies are needed and should employ longitudinal sampling, single-cell RNA-sequencing, or in vitro co-cultures to experimentally confirm how microglia and peripheral immune cells influence OPC differentiation. Second, the use of elderly control subjects introduces an age-related inflammatory baseline that may underestimate the net inflammatory changes seen in EAD. While our groups are age-matched, our controls subjects are around 80-year-old and it is known that inflammation increases with age (Andronie-Cioara et al., 2023; Sparkman & Johnson, 2008). To further study the inflammatory aspect and peripheral infiltration, younger subjects could be added as low-inflammation controls. Finally, the extent to which late stage remyelination attempt could translate into functional benefits remains unknown. Additional mechanistic studies, possibly including advanced imaging or behavioral correlations in animal models, could clarify how changes in the inflammatory milieu impact cognitive outcomes.

Therapeutically, our data support the potential of targeting inflammation in a stage-specific manner. Modulating microglial reactivity or peripheral immune infiltration in early AD might protect OLs from excessive damage as previously reported (Arfaei et al., 2024). Conversely, promoting OPC maturation in later stages may prove beneficial if coupled with strategies to sustain neuronal health. More broadly, the idea of temporal immune modulation, either by decreased early inflammation or enhancing late-stage regenerative responses, deserves further investigation.

In conclusion, our data support a model in which early-stage hyperinflammation leads to OL damage, followed by OL recovery attempts when inflammation subsides. Targeting these stage-specific pathways could offer promising avenues for AD therapy.

## Acknowledgements

We would like to thank Pia Lovero and Maria Surini for technical assistance. We would like to acknowledge the iGE3 Genomics platform and Mylène Docquier for their contribution to the mRNA sequencing. The graphical abstract was made using BioRender. This work was supported by the Swiss National Science Foundation (grant 310030_212322).

## Author contribution

A.M.B.: investigation, formal analyses, writing – original draft. Q.A. formal analyses, writing – review and editing. K.C.: conceptualization, supervised wet lab experiments, writing – review and editing. L.A: writing – review and editing. A.T., L.G. and E.V.: investigation. S.T. and P.M.: conceptualization and writing – review and editing. B.B.T. conceptualization, supervision, funding acquisition and writing – review and editing.

