## Supplementary material for "Non-linear microglial, inflammatory and oligodendrocyte dynamics across stages of Alzheimer’s disease": Supp

### Supplemental data

| Group | Subject | Age (years) | Braak stage | Sex | Post-mortem delay (minutes) |
|---|---|---|---|---|---|
| CT | **1** | 82 | 2 | Male | 355 |
| | **2** | 95 | 3 | Female | 260 |
| | **3** | 87 | 3 | Male | 380 |
| | **4** | 101 | 3 | Female | 360 |
| | **5** | 91 | 3 | Female | 570 |
| | **6** | 84 | 3 | Female | 470 |
| EAD | **7** | 102 | 4 | Female | 310 |
| | **8** | 78 | 4 | Female | 180 |
| | **9** | 96 | 4 | Female | 240 |
| | **10** | 87 | 4 | Female | 240 |
| | **11** | 77 | 4 | Female | 190 |
| | **12** | 79 | 4 | Male | 390 |
| | **13** | 88 | 4 | Female | 600 |
| | **14** | 87 | 4 | Female | 205 |
| | **15** | 86 | 4 | Male | 290 |
| | **16** | 89 | 4 | Male | 355 |
| | **17** | 86 | 4 | Male | 225 |
| | **18** | 76 | 4 | Female | 300 |
| LAD | **19** | 88 | 5 | Female | 405 |
| | **20** | 81 | 6 | Female | 165 |
| | **21** | 81 | 6 | Female | 315 |
| | **22** | 87 | 6 | Female | 480 |
| | **23** | 86 | 6 | Female | 315 |
| | **24** | 76 | 6 | Male | 310 |
| | **25** | 85 | 6 | Female | 315 |
| | **26** | 95 | 6 | Female | 395 |
| | **27** | 81 | 6 | Female | 330 |
| | **28** | 88 | 6 | Female | 379 |
| | **29** | 78 | 6 | Male | 395 |
| | **31** | 75 | 6 | Male | 185 |

**Supplemental Table 1. Fresh-frozen subjects' data table.** There are no differences in age between the CT (mean age of 90 ± 7.16), the EAD (mean age of 85.12 ± 7.77) and the LAD (mean age of 82.36 ± 4.74) groups. ANOVA F(2, 26)=2.628, p=0.0913. There are differences in Braak stages between groups: Kruskal-Wallis $\chi^2$=28.408, p=6.78e-7. Dunn's comparison CT-EAD; Z=-2.18, p.adj=0.029. Dunn's comparison CT-LAD; Z=-5.07, p.adj=1e-6. Dunn's comparison EAD-LAD; Z=-3.55, p.adj=7.6e-4. There are no differences in sex distribution between groups: $\chi^2$=0.238, p=1. There are no differences in post-mortem delay between groups: ANOVA F(2, 27)=2.04, p=0.672.

| Group | Subject | Age (years) | Braak stage | Thal stage | Sex | Post-mortem delay (minutes) |
|---|---|---|---|---|---|---|
| CT | **1** | 86 | 2 | 2 | Female | 2220 |
| | **2** | 76 | 2 | 2 | Male | 1860 |
| | **3** | 76 | 3 | 2 | Female | 3060 |
| | **4** | 85 | 2 | 1 | Female | 4320 |
| | **5** | 73 | 0 | 3 | Female | 1680 |
| | **6** | 74 | 2 | 2 | Male | 2220 |
| | **7** | 82 | 2 | 2 | Male | 2160 |
| | **8** | 84 | 2 | 3 | Female | 1920 |
| | **9** | 71 | 3 | 2 | Female | 1560 |
| | **10** | 91 | 2 | 2 | Male | 2400 |
| | **11** | 86 | 3 | 3 | Male | 1740 |
| | **12** | 86 | 3 | 3 | Male | 1740 |
| | **13** | 87 | 0 | 0 | Male | 1500 |
| | **14** | 87 | 2 | 0 | Male | 2460 |
| | **15** | 79 | 3 | 0 | Male | 1440 |
| | **16** | 89 | 2 | 0 | Female | 2640 |
| | **17** | 51 | 1 | 0 | Male | 5040 |
| | **18** | 89 | 2 | 0 | Female | 2640 |
| | **19** | 82 | 2 | 0 | Female | 2700 |
| | **20** | 69 | 3 | 0 | Female | 5760 |
| | **21** | 79 | 2 | 0 | Male | 5040 |
| | **22** | 89 | 2 | 0 | Female | 1200 |
| | **23** | 54 | 0 | 0 | Female | 1260 |
| | **24** | 85 | 3 | 0 | Male | 1800 |
| | **25** | 84 | 3 | 0 | Female | 3600 |
| EAD | **26** | 87 | 5 | 2 | Female | 5760 |
| | **27** | 86 | 4 | 3 | Male | 2160 |
| | **28** | 93 | 4 | 2 | Female | 2760 |
| | **29** | 92 | 4 | 4 | Female | 4140 |
| | **30** | 91 | 4 | 2 | Female | 2760 |
| | **31** | 90 | 4 | 2 | Female | 300 |
| | **32** | 78 | Unknown | Unknown | Female | 840 |
| | **33** | 79 | 4 | 4 | Male | 2880 |
| | **34** | 82 | 5 | 2 | Male | 4260 |
| LAD | **36** | 91 | 6 | 4 | Male | 1260 |
| | **37** | 90 | 5 | 4 | Male | 2160 |
| | **38** | 77 | 6 | 3 | Male | 1560 |
| | **39** | 92 | 5 | 4 | Male | 1200 |
| | **40** | 91 | 5 | 4 | Female | 1200 |
| | **41** | 88 | 6 | 4 | Female | 1140 |
| | **42** | 93 | 5 | 3 | Male | 2940 |
| | **43** | 91 | 5 | 3 | Male | Unknown |
| | **44** | 74 | 5 | 3 | Female | 3420 |
| | **45** | 85 | 5 | 3 | Female | 3720 |
| | **46** | 75 | 5 | 3 | Male | 4560 |

**Supplemental Table 2. FFPE subjects' data table.** There are no differences in age between the CT (mean age of 79.76 ± 10.2), the EAD (mean age of 86.44 ± 5.64) and the LAD (mean age of 86.09 ± 7.26) groups. There are differences in Braak stages between groups: Kruskal-Wallis $\chi^2$=34.98, p=2.5e-8. Dunn's comparison CT-EAD; Z=-3.43, p.adj=0.018. Dunn's comparison CT-LAD; Z=-5.57, p.adj=7.73e-8. Dunn's comparison EAD-LAD; Z=-1.33, p.adj=0.18. There are differences in Thal stages between groups: Kruska-Wallis $\chi^2$= 21.749, p=1.89e-5. Dunn's comparison CT-EAD; Z=-2.41, p.adj=0.032. Dunn's comparison CT-LAD; Z=-4.49, p.adj=2.2e-5. Dunn's comparison EAD-LAD; Z=-1.47, p.adj=0.14. There are no differences in sex distribution between groups: $\chi^2$=5.42, p=0.2344. There are no differences in post-mortem delay between groups: ANOVA F(2, 41)=0.402, p=0.672.

| EAD vs. CT | LAD vs. CT | LAD vs. EAD | Overlapping EAD vs. CT | Overlapping EAD vs. CT and LAD vs. EAD | Overlapping LAD vs. CT and LAD vs. EAD | All overlapping |
|---|---|---|---|---|---|---|
| TLR7, MEF2A, NINJ2, CNP, S100B, GRIN3A, CCR5, B2M, CABLES1, ANKH, PSMB8 | AKT3, SHC3, SMAD3, FCGR3A/B, GPNMB, GRN, RBX1, UNC79, SCP2, SUGT1, PPM1L, GNB1, ATP6V1G2, TLN2, C1R, TMEM47, PRKAG1, HLA-A, SLC6A1, HAMP, FGF2, KCTD12, SYNGR1, CNTNAP1, CD109, TRIM46, BECN1, DNM3, FCGR1A/B, HSPA1A, ATP2B2, PIAS1, SLC38A2, ENO2, CD44, GNAI1, TBX21, AMIGO2, KITLG, FPR1, TBP, ABCA1, FAM30A | YWHAG, PPP2R2B, ATG5, TIE1, PDGFRA, JUN, CUL2, PPT1, IFNAR1, FABP5, CX3CL1, DRD1, MBP, ADRB1, P2RY11, THY1, HADH, ERBB3, VIM, KLHL12, ANK2, PIK3R1, FIG4, MAL2, BCAS1, RORB, CDK5R1, SLC9B2, TSC1, HDAC2, LAMTOR3, CCNA1, LAMP2, PDPK1, DLX1, TJP1, MERTK, FGF13, GABRA5, ATP2A2, PTH2R, CNTF, SLC46A1, PFKP, SERPINE2, MAP2K4, IL1A, SCN1A, ADORA2B, LARS1, QDPR, PLAA, ORC4, SLC8A1, GUSB, CTSF, LAMP1, PKD2, GRK3, ABCG2, GSN, CTNNB1, UBE2D2, AMIGO1, PIK3CB, MYRF, TICAM1, CAMK2G, INPP5F, E2F3, TIAM1, MYORG, AKT1, EPB41L2, BTRC, LSR, BDNF, DIAPH1, KRAS, RASGRP1, CLDN5, XPNPEP1, HSD17B4, KCND1, TUSC3, SEC23A, NOSTRIN, TRIM26, B4GALT6, RHOB, NRCAM, SALL1, SCD5, SOX10, UBE3C, RAD23B, OLIG1, B4GALT5, ITM2A, CYFIP2, SLC24A3, RIMS1, ARPC2, GABRB3, GLS, NCKAP1, PLXNB3, GRIA1, PTX3 | PTK2, BACE1, PRKCQ, PLLP, GAS7, TFEC, PRKAB1, NOS2, CEP68 | ARG2, MAG, CUL1, GAL3ST1, ALDH1L1, TMEM88B, TRIM45, SORL1, BIN1, AARS1, OLIG2, FTH1, PMP22 | SYN2, RLIM, NCAM2, UBQLN2, XK, RTN4, RANBP9, MAPK1, CSNK2A1/3, DYNC1LI1, ROCK2, PRKCZ, DLAT, CLTA, ESAM, GNG2, MYO5A, P2RX7, ASNS, MBTPS1, MAP2K1, MAD2L1, ADAM22, CAMK2A, NSF, EMCN, EPB41L3, VDAC3, ATP6V1H, AP3M2, SLC9A6, CACNA1B, SYT4, GAD1, MAPK14, MAPK10, SOX9, GABRA3, ATP6V1C1, GRM7, GSK3B, GPM6A, YWHAQ, SYBU, CASP3, PTPN1, MAPK9, YWHAH, ELOC, ZPR1, TMSB4X, C1QBP, CYP2U1, GRIK2, ANAPC1, DCLK1, SLC25A4, ATP6V1A, NDUFB5, APOE, NF1, BRAF, GNG3, SYN1, ATP6V0D1, CSF1, TIMP2, ENTPD2, ATP5PB, UNC80, NLRP1, CACNA2D1, ACSL4, WASF1, ERLEC1, CDK7, CHRM3, PRKACB, TADA2B, HCN1, CNR1, NEFL, OLFM3, SPARC, DNAJC5, CHN1, PDHA1, B3GNT5, ACLY, CYP4X1, PRKCI, MTO1, MEAF6, KCNQ3, RAB6B, PLEKHB1, PGAM1, LRRC7, CA2, ATF2, GABRA4, DNM1, PPP3R1, SERINC3, GRIA2, DNAJB9, REEP5, OPA1, CLTC, GPR17, PTK2B, RASGRF1, APLP2, PPP3CB, FCGRT, CALM1, CD47, ATRN, PARP2, SCOC, NRXN1, INA, KCND2, UBE2N, ATP6V1D, SLC12A5, TPI1, CDC27, MAPK8, DLGAP1, ATP1B1, ATP1A1, APP, CHRNB2, AMPH, TRIM3, CD200, ATP8A2, GRIN2A, PSMD6, GPRASP1, ATP6V1G1, SELPLG, UCHL1, SH3GL2, GLRB, YWHAE, GABARAPL1, GABRA1, PPP2CA, YWHAZ, HK1, VDAC1, PAK1, CAMK4, TLR4, ACAT2, IMMT, PSMC6, STXBP1, RTN1, ATR, GAD2, EGFR, OPCML, FBLN5, PSMD14, SCN8A, PDHB, TSPAN7, DLD, S100A10, P2RY1, PRKAR1A, RAB3A, KCNAB1, SUMO1, EGF, ATP6V1B2, KCNV1, DNAJA2, NES, TRIM37, SQSTM1, ACADSB, SOX2, TXN, NAP1L2, NPTN, CDC42, NMNAT2, PDE1A, ARPC5L, TNFRSF21, SYT1, PGK 1.00, GOT1, TANC2, PSMA5, SLC25A12, NWD1, CNTNAP2, ATP5MC3, GRIA3, SNAP25, DNM1L, ABCE1, LGMN, GABRG2, NKIRAS1, EID1, RET, ADCYAP1R1, AXL, CALM2, SKP1, ATP5F1B, ATP6V1E1, GRB2, GFAP, GDPD2, C4A/B, DNAJC11 | NDUFA10, SEC61A2, GPHN |

**Supplemental Table 3. Genes with significantly modulated expression from the Venn diagram in Figure 1.D.** A total of 411 genes had dysregulated expression.

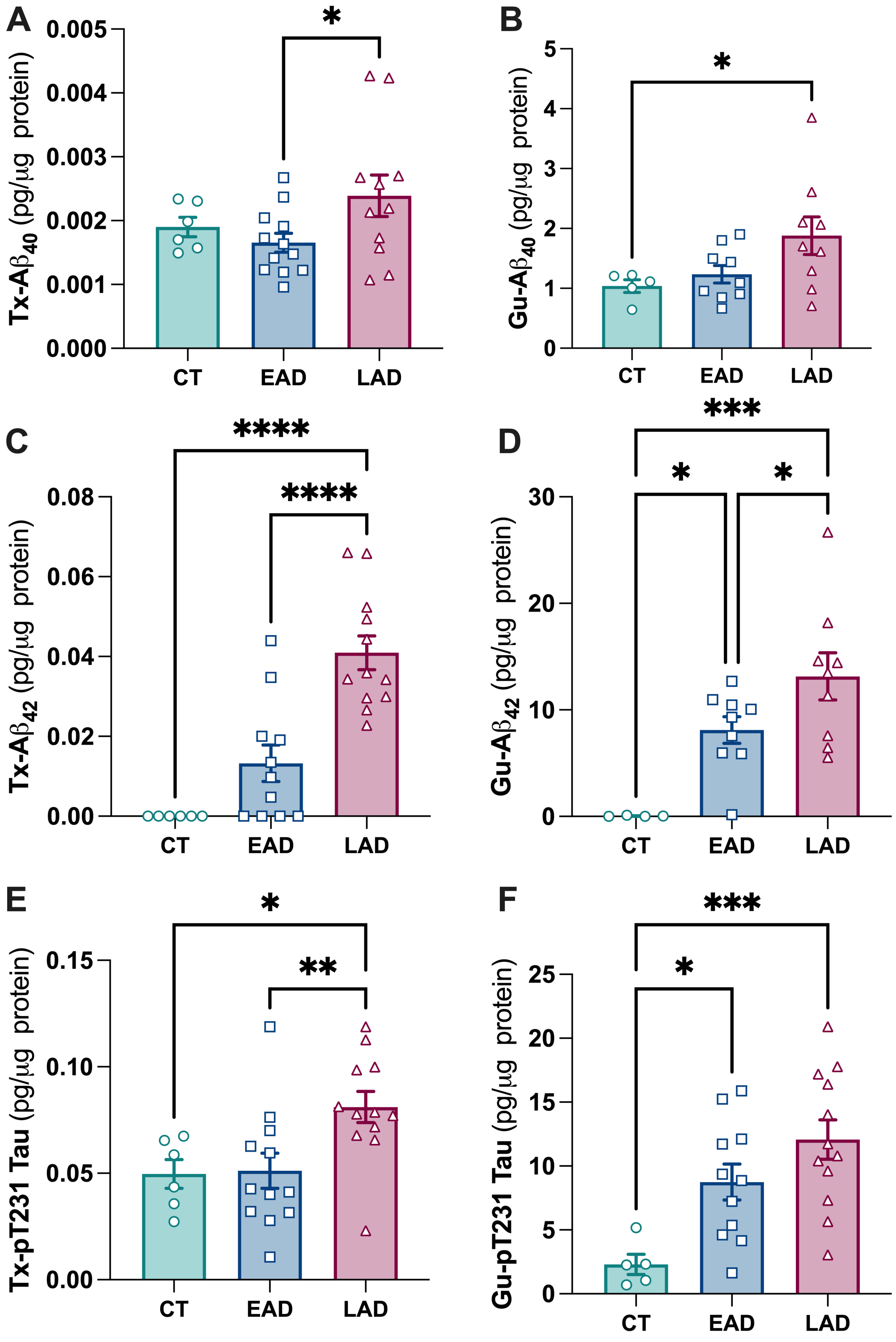


**Supplemental figure 1. Soluble and insoluble protein markers in the different groups. A.** Aβ40 extracted with Triton (Tx) buffer. **B.** Aβ40 extracted with Guanidine (Gu) buffer **C.** Aβ42 extracted with Triton buffer. **D.** Aβ42 extracted with Guanidine buffer. **E.** Phosphorylated Tau extracted with Triton buffer. **F.** Phosphorylated Tau extracted with Guanidine buffer. Statistics were done with one-way ANOVA and uncorrected Fisher's LSD post hoc test. CT: Controls, EAD: Early AD, LAD: Late AD.

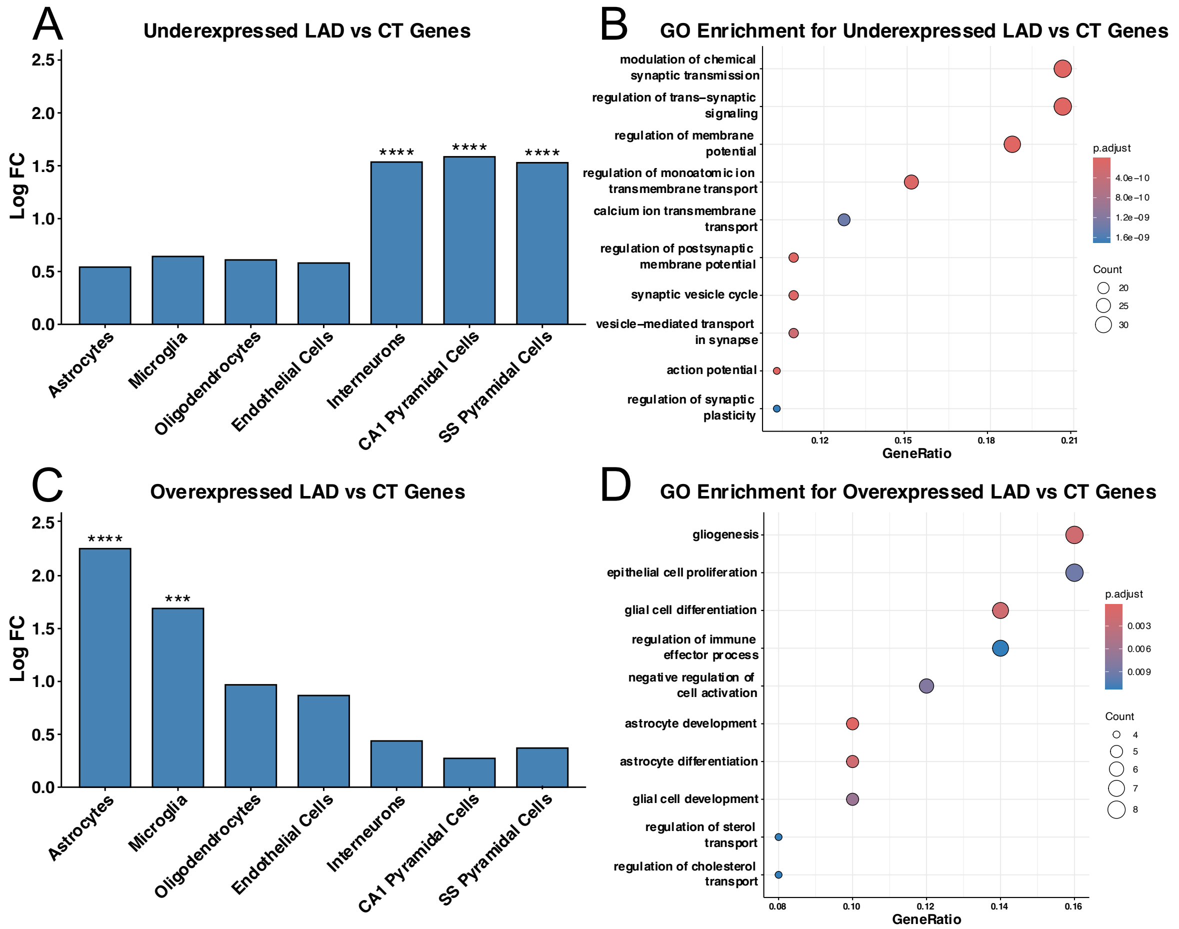


**Supplemental figure 2. Cellular and pathway enrichment in gene changes in LAD versus CT. A.** Cellular deconvolution show that the genes with downregulated expression are expressed by neurons. **B.** Enrichment of genes identified in A reveal neuronal pathways. **C.** Cellular deconvolution show that the genes with upregulated expression are expressed by astrocytes and microglia. **D.** Enrichment of genes identified in C reveal immune pathways such as glial cells regulation, proliferation, astrocyte development, regulation of immune processes. CT: Controls, LAD: Late AD. *** for $p < 0.001$, **** for p<1e-4.
